# Neuroanatomical features reveal accelerated brain age in alcohol use disorder

**DOI:** 10.1101/2025.09.30.679671

**Authors:** Chella Kamarajan, Babak A. Ardekani, Ashwini K. Pandey, Jacquelyn L. Meyers, David B. Chorlian, Sivan Kinreich, Gayathri Pandey, Stacey Saenz deViteri, Christian Richard, Arjun Bingly, Weipeng Kuang, Bernice Porjesz

**Affiliations:** Henri Begleiter Neurodynamics Laboratory, Department of Psychiatry and Behavioral Science, SUNY Downstate Health Sciences University, Brooklyn, NY 11203, USA; Center for Biomedical Imaging and Neuromodulation, Nathan Kline Institute for Psychiatric Research, Orangeburg, NY 10962, USA; Department of Psychiatry, NYU School of Medicine, New York, NY 10016, USA

**Keywords:** Brain age, alcohol use disorder, cortical thickness, brain volume, neuropsychological performance, brain health

## Abstract

**Background/Objectives:** Brain age is a novel measure to characterize the integrity of neurocognitive functioning and brain health in various psychiatric and neurological disorders. Although there is a literature suggesting premature aging of the brain in individuals with alcohol use disorder (AUD), studies directly examining brain age are rare. Therefore, the current study was designed to estimate brain age in AUD individuals using brain morphological features, such as cortical thickness and brain volume.

**Methods:** The sample included a group of 30 adult males with a history of AUD but maintaining abstinence and a group of 30 male controls without any history of AUD. Brain age was computed using an XGBoost regression model with 187 brain morphological features of cortical thickness and brain volume as predictors. An exploratory correlational analysis between brain age measures and features of neuropsychological performance, impulsivity, and alcohol consumption was also performed.

**Results:** Findings revealed that AUD individuals showed an increase of 1.70 years in their brain age relative to their chronological age. Further, in the AUD group, higher brain age was significantly associated with poor executive functioning, while a larger gap between brain age and actual age was associated with lower non-planning impulsivity and later age of onset for regular drinking in those with AUD.

**Conclusions:** AUD individuals manifested accelerated brain aging, possibly representing compromised brain health. Brain age measures were found to be associated with some of the measures of neurocognition, impulsivity, and alcohol consumption. These findings may have important implications for the early identification, prevention, and treatment of AUD. However, future studies with larger sample sizes are needed to confirm these preliminary findings.

## 1. Introduction

Prediction of brain age relative to one’s chronological age can serve as a useful biomarker of brain health [1–3]. Brain age estimation using neural measures has been successfully used as a measure of neurobiological aging and cognitive decline/lag [4–6] in clinical populations [1,3,7,8]. Brain age indices computed from structural and functional brain measures, especially using brain morphological features, are increasingly being used to understand brain integrity, neurodevelopment, and neuropsychiatric disorders [3,9–11]. There is substantial evidence that substance use disorders (SUD) can lead to cognitive impairment [12–14], and recent studies have reported that individuals with SUD have manifested accelerated brain age [15,16]. Particularly, although individuals with alcohol use disorder (AUD) often manifest aberrations in both structural and functional brain measures [17,18], studies on brain age in AUD are rare [19], which limits our understanding of the impact of chronic use of alcohol on brain health. Further, it is less clear from the available reports whether brain age is associated with other measures of cognition and behavior, such as neuropsychological performance and impulsivity. In order to provide further information about the impact of AUD on brain health, we examine brain age in individuals with a history of AUD, in addition to its association with various measures of neurocognition, impulsivity, and alcohol consumption.

Morphological measures such as brain volume and cortical thickness have been primarily used to estimate brain age [1,7,20]. The quantified value by subtracting the computed brain age from the chronological age of a person has been variously termed as brain age gap (BAG), brain age delta (BAD), brain-predicted age difference (Brain-PAD), and brain age gap estimation (BrainAGE) [21]. A positive brain age gap indicates accelerated brain aging, while a negative gap suggests slower aging compared to other individuals in the sample [1]. Brain age measures are typically calculated by first modeling the relationship between selected brain measurements and chronological age within a healthy (or control) sample to establish a baseline [3]. This trained predictive model, typically using a machine learning technique, is then applied to a sample of interest (usually the clinical group) to predict brain age at the individual level [3]. Recently, there has been a growing effort to develop and evaluate methods for measuring brain age using machine learning models. A growing number of recent studies also utilize the popular and efficient ensemble algorithm, XGBoost [22], due to its superior predictive power and performance over other machine learning techniques [9,10,16,23,24].

Earlier studies indicated that chronic alcohol consumption led to “premature aging” [25,26], especially in males [27]. According to this hypothesis, individuals with AUD manifest atrophic changes in their brains [28], which resemble those of age-related brain changes in older individuals [28,29]. Notably, for instance, Pfefferbaum et al. [30] reported that volume loss in both gray and white matter, along with increased volume of cerebrospinal fluid (CSF), was pronounced in individuals with chronic AUD, after correcting for chronological age. Specifically, the volume loss was found to be more pronounced in the frontal cortices and hippocampal regions [18,31,32]. Similar to volumetric changes, individuals with AUD have been shown to have thinner cortices across widespread areas of the brain [33]. A recent review of large-scale neuroimaging studies revealed that both low-to-moderate consumption and heavy consumption of alcohol were associated with smaller volume and cortical thickness in frontal, temporal, and parietal cortices, coupled with volume reductions in insula, cingulate, and subcortical structures [34]. These structural changes in AUD individuals often lead to neurocognitive deficits [35], such as poor executive functioning [36,37] and impaired memory [37,38]. Evidence suggested a predominant impairment of prefrontal cortices and hippocampal regions that underlie these functional deficits in AUD individuals [32,39,40]. Further, accumulating evidence suggests that age-related cognitive decline was accelerated in individuals with AUD [30,41]. Thus, chronic alcohol use can accelerate neurodegeneration, leading to increased risk for premature aging, cognitive decline, and dementia [28,42,43].

Since AUD remains a leading contributor to preventable and premature mortality in the United States [44] and worldwide [45], elucidation of brain age profiles in individuals with a history of AUD may provide a better understanding of and treatment for AUD. Therefore, the current study is an attempt to estimate brain age in a group of male adults with a lifetime diagnosis of AUD, in comparison to a group of healthy individuals without a history of AUD, following a recent study by Rutherford et al. [46]. As most of the recent studies on brain age have used MRI-based neuroanatomical features to predict brain age in several disorders [1,6–9,46–49], the present study has also utilized these measures, specifically brain volumes and cortical thickness. Since individuals with a history of AUD have also been shown to manifest impaired neuropsychological functioning [40,50–55] and heightened impulsivity [40,56–60], the current study has also attempted to investigate the association of brain age with neuropsychological scores (e.g., executive function and working memory) and impulsivity scores. We hypothesized that individuals with a history of AUD would manifest accelerated brain age compared to control individuals without AUD. It was expected that the findings from the present study would enhance our understanding of brain aging in individuals with AUD, as well as of the associations among brain age, neurocognition, and impulsivity.

## 2. Materials and Methods

### 2.1. Participants

The sample comprised a total of 60 male participants (age range = 19.75–51.08 years), in which 30 individuals had a lifetime diagnosis of DSM-IV alcohol dependence (AUD group; M_age =_ 41.25; SD_age =_ 7.20; age range = 25.58–51.08 years) and 30 individuals did not have any AUD diagnosis (CTL group; M_age =_ 27.24; SD_age =_ 4.78; age range = 19.75–38.08 years) [see **Table 1**]. The AUD individuals were recruited from alcoholism treatment centers in and around NYC after they had been detoxified and did not have any withdrawal or other major AUD symptoms during the time of scanning. CTL individuals were recruited through advertisements and screened to exclude any personal and/or family history of major medical, psychiatric, or substance-related disorders. A detailed description is provided in our previous publications [32,61]. An abbreviated version of the semi-structured assessment for the genetics of alcoholism (SSAGA) [62], a polydiagnostic clinical interview, was administered to assess alcohol/substance use and related disorders. Participants were instructed to abstain from alcohol and other substances for at least 5 days before the scanning. Standard MRI safety protocols and exclusion criteria (implants, tattoos, cosmetics, claustrophobia, etc.) were followed to ensure subjects’ safety and data quality. Individuals with hearing/visual impairment, a history of head injury, or moderate and severe cognitive deficits (<21) on mini-mental state examination (MMSE) [63] were also excluded from the study. All participants provided informed consent, and the Institutional Review Board approved the study protocols.

**Table 1.** Comparison of multidomain features (age, alcohol/drug consumption, impulsivity, and neurocognition) between the AUD and control participants.

| Variables | AUD |  |  | Control |  |  | t | p |
| --- | --- | --- | --- | --- | --- | --- | --- | --- |
|  | N<sup>a</sup> | Mean | SD | N<sup>a</sup> | Mean | SD |  |  |
| Age (in years) | 30 | 41.25 | 7.20 | 30 | 27.24 | 4.78 | 8.78 | <0.001*** |
| Alcohol: Age of onset (regular use)<sup>b</sup> | 30 | 15.77 | 2.58 | 12 | 20.5 | 3.8 | -4.67 | <0.001*** |
| Alcohol: Quantity/day (heavy use period)<sup>c</sup> | 30 | 11 | 7.66 | 12 | 3 | 1.6 | 5.58 | <0.001*** |
| Alcohol: Frequency/month (heavy use period)<sup>c</sup> | 30 | 20 | 9.01 | 12 | 4 | 4.09 | 7.97 | <0.001*** |
| Alcohol: Quantity/day (last 6 months)<sup>c</sup> | 30 | 3 | 6.61 | 18 | 3 | 1.98 | 0.045 | NS |
| Alcohol: Frequency/month (last 6 months)<sup>c</sup> | 30 | 4 | 8.02 | 18 | 3 | 3.62 | 0.6 | NS |
| Alcohol: Length of Abstinence (in days)<sup>d</sup> | 30 | 672.93 | 844.94 | 18 | 57 | 149.76 | 3.89 | <0.001*** |
| Tobacco: Quantity/day (last 6 months)<sup>c</sup> | 20 | 10 | 5.8 | 6 | 2 | 1.63 | 5.19 | <0.001*** |
| Tobacco: Frequency/month (last 6 months)<sup>c</sup> | 20 | 28 | 4.83 | 6 | 14 | 13.82 | 2.47 | NS |
| Marijuana: Frequency (last 6 months)<sup>c</sup> | 10 | 99 | 91.38 | 4 | 19 | 27.61 | 2.58 | <0.03* |
| BIS Non-planning | 28 | 24.57 | 5.25 | 30 | 19.80 | 4.61 | 3.69 | <0.001*** |
| BIS Motor impulsivity | 27 | 25.15 | 5.24 | 30 | 19.30 | 3.28 | 5.10 | <0.001*** |
| BIS Attentional impulsivity | 28 | 15.79 | 4.13 | 30 | 12.57 | 3.19 | 3.33 | 0.0015** |
| BIS Total impulsivity | 27 | 65.44 | 11.51 | 30 | 51.67 | 8.60 | 5.15 | <0.001*** |
| TOL_MovMade | 25 | 93.04 | 17.02 | 29 | 85.83 | 6.66 | 2.10 | 0.0402* |
| TOL_ExcMovMade | 25 | 15.04 | 17.02 | 29 | 7.83 | 6.66 | 2.10 | 0.0402* |
| TOL_PctAbvOpt | 25 | 19.28 | 21.82 | 29 | 10.04 | 8.54 | 2.10 | 0.0402* |
| TOL_AvgPicTime | 25 | 3.09 | 1.09 | 29 | 2.81 | .96 | 1.01 | 0.3167 |
| TOL_AvgTotTime | 25 | 5.16 | 1.58 | 29 | 4.72 | 1.64 | 1.00 | 0.3199 |
| TOL_TotTrlTime | 25 | 482.60 | 178.18 | 29 | 404.24 | 139.05 | 1.81 | 0.0755 |
| TOL_AvgTrlTime | 25 | 22.98 | 8.48 | 29 | 19.25 | 6.62 | 1.81 | 0.0755 |
| VST_TotCor_Fw | 25 | 7.00 | 2.58 | 29 | 10.21 | 2.78 | -4.37 | <0.001*** |
| VST_Span_Fw | 25 | 5.44 | 1.33 | 29 | 6.83 | 1.36 | -3.78 | <0.001*** |
| VST_TotAvgTime_Fw | 25 | 26.72 | 9.13 | 29 | 28.31 | 10.53 | -0.59 | 0.5591 |
| VST_TotCorAvgTime_Fw | 25 | 38.15 | 12.39 | 29 | 32.48 | 8.07 | 2.02 | 0.0490* |
| VST_TotCor_Bw | 26 | 6.31 | 3.02 | 29 | 8.31 | 1.87 | -2.99 | 0.0042** |
| VST_Span_Bw | 26 | 4.65 | 1.44 | 29 | 5.52 | .95 | -2.65 | 0.0106* |
| VST_TotAvgTime_Bw | 26 | 17.72 | 9.39 | 29 | 17.79 | 10.01 | -0.03 | 0.9791 |
| VST_TotCorAvgTime_Bw | 26 | 28.93 | 13.92 | 29 | 27.16 | 10.66 | 0.53 | 0.5963 |

<sup>a</sup> Note that all participants were assessed, but the data of participants who did not endorse specific items were omitted from the analysis here.
<sup>b</sup> Drink one day/per month for 6 months or more.
<sup>c</sup> The mean values of quantity and frequency values were rounded to the nearest whole numbers.
<sup>d</sup> Range (5–4228 days).
\* $p < 0.05$ (statistically significant at 5% level)
\*\* $p < 0.01$ (statistically significant at 1% level)
\*\*\* $p < 0.001$ (statistically significant at 0.1% level)

### 2.2. MRI Data Acquisition and Image Processing

Details on MRI data acquisition are available in our previous publication [32]. The scanning was done using Siemens 3T Tim *Trio*. A high-resolution three-dimensional T1-weighted magnetization-prepared rapid gradient-echo (MPRAGE) image was collected with the following parameters: TR = 2500 ms, TE = 3.5 ms, TI = 1200 ms, flip angle = 8°, matrix size = 256×256×192, and voxel size = 1×1×1 mm^3^. Standard procedures and protocols of MRI scanning were followed. A total of 187 brain morphological features [see **Table A1** in the *Appendix section*] were extracted for the prediction of brain age, based on the previous study by Rutherford et al. [46]. We used the *sMRIPprep* (version 0.12.2) workflow [64], which is part of a Python-based image analysis platform *Nipype* (version 1.8.6) [65], for preprocessing anatomical data from the T1-weighted (T1w) images. Complete details of the software, processing, processing pipeline, and documentation are available at https://www.nipreps.org/smriprep/. The preprocessing pipeline included the following steps. First, the T1-weighted (T1w) image was corrected for intensity non-uniformity (INU) with *N4BiasFieldCorrection* [66], distributed with *ANTs* (version 2.3.3) [67] and used as T1w-reference throughout the workflow. The T1w-reference was then skull-stripped with a Nipype implementation of the *antsBrainExtraction.sh* workflow (from ANTs), using *OASIS30ANTs* as the target template. Brain tissue segmentation of cerebrospinal fluid (CSF), white matter (WM), and gray matter (GM) was performed on the brain-extracted T1w using *fast* from FSL (version 6.0.5.1:57b01774) [68]. Brain surfaces were reconstructed using *recon-all* from FreeSurfer (version 7.3.2) [69], and the brain mask estimated previously was refined with a custom variation of the method to reconcile ANTs-derived and FreeSurfer-derived segmentations of the cortical gray matter of *Mindboggle* [70].

Cortical thickness values of the Destrieux parcellation [71] containing 74 regions and a mean value for each hemisphere, along with volumetric measures for 37 brain structures, were extracted from the Freesurfer output directories. All 187 neuroanatomical features (150 on cortical thickness and 37 on brain volumes) were used in the brain age prediction model.

### 2.3. Brain Age Prediction Model

We implemented an *XGBoost* [22] (stands for *eXtreme Gradient Boosting*) as several recent studies have used this algorithm to predict brain age using MRI [72–77] as well as EEG measures [78]. The details of the algorithm and computational steps are available in the work by de Lange et al. [9], wherein the algorithms from the native XGBoost [22] and Scikit-learn [79] have been implemented for computing brain age. As mentioned earlier, 187 anatomical features (i.e., 150 features of bilateral cortical thickness and 37 features of brain volumes) [see **Table A1** in the *Appendix section*] served as the predictor variables against chronological age as the response or outcome variable in the XGBoost regression model. We tested the prediction model using data from the CTL group and tested the model using data from the AUD group, as it is usually done in previous brain age studies on clinical groups (see Seitz-Holland et al. [3] for a review). Predicted brain age was subjected to an age-bias correction procedure [9,80]. While training the model, the values of the predictor variables (i.e., the neuroanatomical features) were scaled using the *robust scaler* [81] from the *scikit-learn* library [79], which removes the median and scales the data according to the quantile range [9]. Hyperparameter tuning was implemented to obtain the best model parameters to use in the training phase, based on the previous work [9]. We tuned the model parameters using the inner 3-fold and outer 5-fold cross-validation procedure, following the work by de Lange et al. [9]. The *SearchTerm* space parameters in the model were: *max_depth* = range(1, 11, 2), *n_estimators* = range(50, 400, 50), *learning_rate* = [0.001, 0.01, 0.1, 0.2].

It is essential to acknowledge that predicted brain age can be biased, as initial estimates tend to overpredict in younger individuals and underpredict in older individuals before any correction procedures are applied [80], and therefore, bias-correction methods are recommended to correct the initially predicted brain age values. In the current study, we have used the correction procedure developed in a recent work by de Lange et al. [9], who demonstrated brain age prediction using neuroanatomical features and the XGBoost model. According to this procedure, a correction is applied to the predictions by first fitting *Y = α × Ω + β*, where Y is the modeled predicted age as a function of chronological age (Ω), and α and β represent the slope and intercept [9]. The derived values of α and β are then used to correct predicted age with *Corrected Predicted Age = Predicted Age + [Ω − (α × Ω + β)].* The brain age gap, a measure of lag or difference between the chronological age and the predicted brain age [9,82], was also computed. The model performance was measured using the correlation coefficient (r), r-square (R^2^), Mean Absolute Error (MAE), and Root Mean Squared Error (RMSE). In this study, the correlation coefficient (r) between the chronological age and corrected predicted age served as the primary model performance. A higher r-value (regardless of direction) indicated better performance. R^2^, which is not the square of r in this case, is a measure of the predictive power of the regression in terms of how much variation is explained by the regression, and therefore higher R^2^ represents better performance. The MAE represents the average of the absolute value of each residual, and a lower MAE will imply better performance. On the other hand, the RMSE is computed as the square root of the average of squared errors, where the errors are squared before they are averaged. Similar to MAE, higher RMSE signifies lower performance. The MAE and the RMSE together are useful to estimate the variation in the errors in a set of predictions. The RMSE is larger or equal to the MAE, and the difference between the two measures reflects the magnitude of the variance in the individual errors in the sample.

### 2.4. Neuropsychological Assessment

Participants were administered two computerized tests from the Colorado Assessment Tests for cognitive and neuropsychological assessment [83], namely, the Tower of London Test (TOL) [84], and the visual span test (VST) [85]. Details are available in our previous publication [86]. The TOL assesses the planning and problem-solving ability of the executive functions. In this test, participants were asked to solve a set of puzzles with graded difficulty levels by arranging a specific number of colored beads one at a time from a starting position to the desired goal position in as few moves as possible. The test consisted of 3 puzzle types, with 3, 4, and 5 colored beads placed on the same number of pegs, with 7 problems/trials per type and a total of 21 trials. The following performance measures from the sum total of all puzzle types were used in the analysis: (i) actual moves made (MovMade), (ii) excess moves, which are the additional moves beyond the minimum moves required to solve the puzzle (“ExcMovMade”); (iii) percentage of excess moves (PctAbvOpt), (iv) average pickup time, which is the initial thinking/planning time spent until picking up the first bead to solve the puzzle (“AvgPicTime”); (v) average total time, which is the total thinking/planning time to solve the problem in each puzzle type (“AvgTotTime”); (vi) total trial time, which is the total performance/execution time spent on all trials within each puzzle type (“TotTrlTime”); and (vii) average trial time, which is the mean performance/execution time across trials per puzzle type (“AvgTrlTime”).

The VST measured the visuospatial memory span from the forward condition and working memory from the backward condition. In this test, a set of randomly arranged squares, ranging from 2 to 8 squares per trial, flashed in a predetermined sequence depending on the span level being assessed. Each span level was administered twice, with a total of 14 trials in each condition. During the forward condition, subjects were required to repeat the sequence in the same order by clicking on the squares using a computer mouse. In the backward condition, subjects were required to repeat the sequence in the reverse order (starting from the last square). The following performance measures were collected during forward and backward conditions: (i) total number of correctly performed trials (“TotCor_Fw” and “TotCor_Bw”); (ii) maximum span or sequence-length achieved (“Span_Fw” and “Span_Bw”); (iii) total average time, which is the sum of mean time-taken across all trials performed (“TotAvgTime_Fw” and “TotAvgTime_Bw”); and (iv) total correct average time, which is the sum of mean time-taken across all trials correctly performed (“TotCorAvgTime_Fw” and “TotCorAvgTime_Bw”).

### 2.5. Impulsivity Measures

Barratt Impulsiveness Scale—Version 11 (BIS-11) [87], a 30-item self-administered tool with excellent psychometric properties [88], was used to assess aspects of impulsivity.

Three subscales, viz., motor impulsivity (BIS_MI), non-planning (BIS_NP), and attentional impulsivity (BIS_AI), along with the total score, were included in the correlational analysis with the age measures.

### 2.6. Alcohol Consumption Measures

Four variables related to alcohol consumption, extracted from the modified short form of SSAGA [32,89], were used in correlational analyses with brain age measures: (i) Age of onset for regular drinking; (ii) Number of drinking days per month during regular drinking period; (iii) Number of drinks per day during regular drinking period; and (iv) Maximum number of drinks in a day during the past 6 months.

### 2.7. Statistical Analysis

Statistical analyses were done using the SPSS software package (Version 28.0.1.1). We compared the demographic and clinical characteristics between the AUD and control groups using independent samples t-tests. Pearson correlations were used to find associations among all age variables (i.e., chronological age, corrected brain age, corrected brain age gap) as well as between the age measures and alcohol consumption and neuropsychological variables. Distributions of brain age measures were plotted for visualization.

## 3. Results

### 3.1. Comparison of Multidomain Variables Across the Groups

As shown in **Table 1**, abstinent AUD participants were significantly younger than control participants. AUD participants started drinking earlier than controls. During their past heavy use period, AUD participants drank more in terms of quantity and frequency than control participants who endorsed drinking. However, the quantity and frequency of drinking during the past 6 months were not significantly different between the AUD and control participants. At the time of assessments, the AUD individuals were maintaining abstinence from drinking longer than the control participants who endorsed drinking. Tobacco use was significantly higher among the AUD participants during the past 6 months than among the control participants who endorsed tobacco use, although the frequency of use was not statistically significant. Marijuana use during the past months was also higher among AUD participants compared to the controls.

### 3.2. Brain Age Measures

Both corrected and uncorrected measures of brain age and brain age gap (brain age – chronological age) were computed. In the XGBoost regression model, the optimal model parameters were as follows: n_estimators = 200, max_depth = 9, and learning_rate = 0.01.

The prediction performance values during the training phase in terms of the absolute mean (SD) values for R^2^, MAE, and RMSE are: 0.096 (0.096), 3.814 (1.828), and 4.443 (1.672), respectively. **Figure 1** shows the distribution of the individual values of Corrected Brain Age against Chronological Age (left panels) and Corrected Brain Age Gap (right panels) for the Control group (top panels), AUD group (middle panels), and the full sample (bottom panels). It can be observed that a majority of the AUD individuals show increased brain age (shown by the red dots over the referenced diagonal line of the bottom left panel) as well as increased brain age gap (shown by the red dots over the “zero” reference line of the bottom right panel).

**Figure 1.**
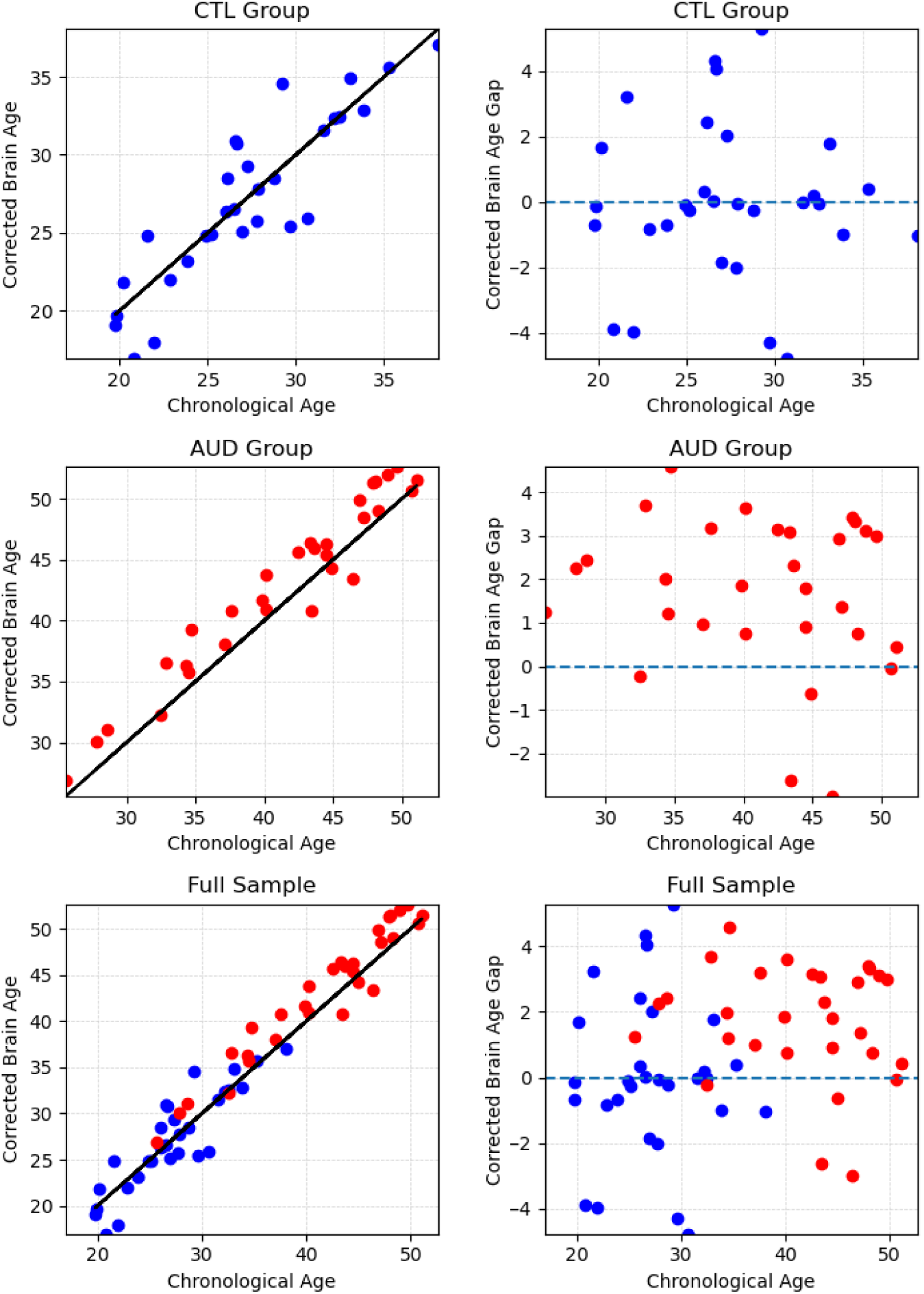
Distribution of the individual values of Corrected Brain Age against Chronological Age (left panels) and Corrected Brain Age Gap against Chronological Age (right panels) for the Control group (top panels), AUD group (bottom panels), and the full sample. The individual values (in blue dots for CTL and in red dots for ALC) above the reference lines in the left panels (brain age) and right panels (brain age gap) represent increased brain age and brain age gap, respectively.

Bar Graphs in **Figure 2** show the mean values for all age measures (Chronological Age, Uncorrected Brain Age, and Corrected Brain Age) [top panel] as well as for the brain age gap measures (Uncorrected Brain Age Gap and Corrected Brain Age Gap) [bottom panel]. It is shown that the Uncorrected Brain Age (29.01 years) in the AUD group was possibly underpredicted by the model (before age-bias correction) against the Chronological Age (41.25 years) during the testing phase. On the other hand, the Corrected Brain Age (42.95 years) was 1.70 years higher than the Chronological Age (41.25 years) in the AUD group, as illustrated in the bar graphs. The control group, which was the training dataset, did not show visible changes in the brain ages. The increased brain age gap in the AUD group, which is 1.70 years, is clearly shown in the bottom panel.

**Figure 2.**
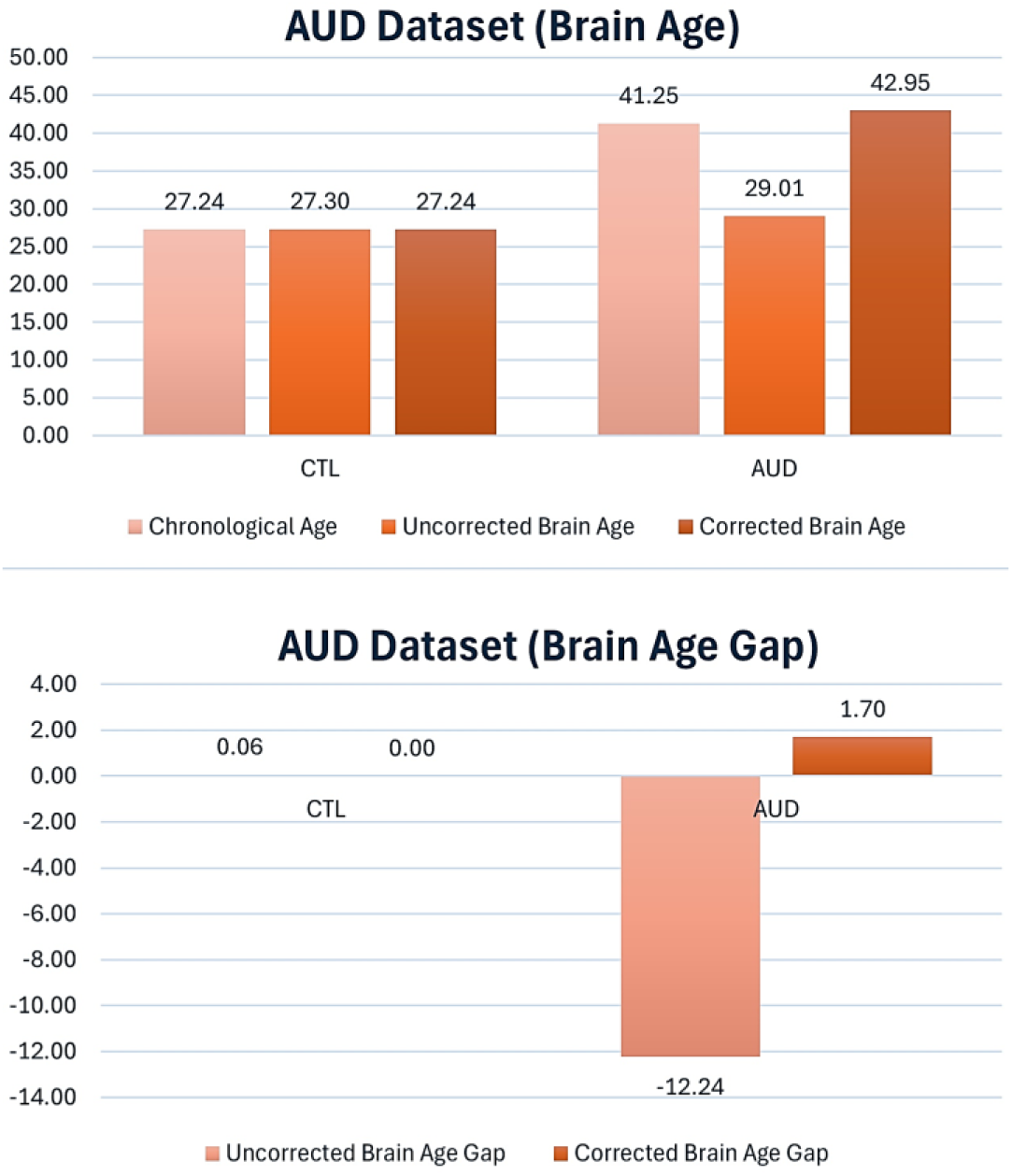
Bar Graphs showing the mean values for Chronological Age, Uncorrected Brain Age, and Corrected Brain Age (top panel) as well as for Uncorrected Brain Age Gap and Corrected Brain Age Gap (bottom panel).

### 3.3. Correlations across Age Measures and Other Variables

#### 3.3.1. Correlations across Different Age Measures

Pearson correlations among the brain age measures in the AUD and CTL groups are shown in **Table 2**. In the Control group, significant correlations were observed between (1) Chronological Age and Corrected Brain Age, (2) Chronological Age and Uncorrected Brain Age Gap, (3) Uncorrected Brain Age and Corrected Brain Age, (4) Uncorrected Brain Age and Uncorrected Brain Age Gap, (5) Uncorrected Brain Age and Corrected Brain Age Gap, (6) Corrected Brain Age and Uncorrected Brain Age Gap, (7) Corrected Brain Age and Corrected Brain Age Gap, and (8) Uncorrected Brain Age Gap and Corrected Brain Age Gap. On the other hand, the AUD group showed significant correlations between (1)

**Table 2.** Pearson correlations among the brain age measures. The correlation values in the lower triangle represent the control group, and those in the upper triangle represent the AUD group. Significant correlations are highlighted in bold font.

|  |  | AUD |  |  |  |  |
| --- | --- | --- | --- | --- | --- | --- |
|  |  | Chronological Age | Uncorrected Brain Age | Corrected Brain Age | Uncorrected Brain Age Gap | Corrected Brain Age Gap |
| CTL | Chronological Age | - | -0.1457 | **0.9691\*\*\*** | **-0.9731\*\*\*** | -0.1502 |
|  | Uncorrected Brain Age | 0.0022 | - | 0.1031 | **0.3696\*** | **1.0000\*\*\*** |
|  | Corrected Brain Age | **0.8903\*\*** | **0.4574\*** | - | **-0.8861\*\*\*** | 0.0985 |
|  | Uncorrected Brain Age Gap | **-0.8900\*\*** | **0.4539\*** | **-0.5847\*\*** | - | **0.3739\*** |
|  | Corrected Brain Age Gap | 0 | **1.0000\*\*** | **0.4555\*** | **0.4559\*** | - |

\* $p < 0.05$ (statistically significant at 5% level)
\*\* $p < 0.01$ (statistically significant at 1% level)
\*\*\* $p < 0.001$ (statistically significant at 0.1% level)

Chronological Age and Corrected Brain Age, (2) Chronological Age and Uncorrected Brain Age Gap, (3) Uncorrected Brain Age and Uncorrected Brain Age Gap, (4) Uncorrected Brain Age and Corrected Brain Age Gap, (5) Corrected Brain Age and Uncorrected Brain Age Gap, and (6) Uncorrected Brain Age Gap and Corrected Brain Age Gap.

#### 3.3.2. Correlations of Age Measures with Neuropsychological Performance

Correlations of specific age measures with neuropsychological scores from the Tower of London (TOL) Test and the Visual Span Test (VST) in AUD and CTL groups are shown in **Table 3**. It is observed that three of the TOL neuropsychological variables related to the number of moves made (i.e., total moves, excess moves, and percentage above optimal moves) showed significant positive correlations with both chronological age and corrected brain age, but only in the AUD group, suggesting poorer performance in executive functioning (planning) with increasing chronological and brain ages. Further, two of the neuropsychological variables related to forward trials in VST (i.e., total correct and maximum span) showed significant negative correlations with chronological age in the AUD group, suggesting poor memory performance with advancing age. On the other hand, the brain age gap was not associated with any of the neuropsychological variables in either group, and there were no significant correlations observed in the CTL group participants across any variables.

**Table 3.** Correlations of specific age measures with neuropsychological scores from the Tower of London (TOL) Test and the Visual Span Test (VST). Note that lower scores in TOL represent better performance, while higher scores in VST indicate better performance. Significant correlations are highlighted in bold font. Correlations with a moderate effect size (r = 0.3 – 0.5), but not statistically significant, have been marked with an omicron superscript (^ο^).

| Neuropsychological Variables<sup>ψ</sup> | AUD |  |  | CTL |  |  |
| --- | --- | --- | --- | --- | --- | --- |
|  | Chronological Age | Brain Age<sup>†</sup> | Brain Age Gap<sup>†</sup> | Chronological Age | Brain Age<sup>†</sup> | Brain Age Gap<sup>†</sup> |
| TOL_MovMade | **0.4909\*** | **0.4924\*** | -0.0156 | 0.0236 | 0.0535 | 0.0691 |
| TOL_ExcMovMade | **0.4909\*** | **0.4924\*** | -0.0156 | 0.0236 | 0.0535 | 0.0691 |
| TOL_PctAbvOpt | **0.4909\*** | **0.4924\*** | -0.0156 | 0.0235 | 0.0535 | 0.0691 |
| TOL_AvgPicTime | 0.0761 | 0.0334 | -0.1986 | 0.1849 | 0.2519 | 0.1828 |
| TOL_AvgTotTime | 0.1555 | 0.0901 | -0.3064$^{\circ}$ | 0.1759 | 0.2296 | 0.1526 |
| TOL_TotTrlTime | 0.3801$^{\circ}$ | 0.3399$^{\circ}$ | -0.2016 | 0.1843 | 0.2374 | 0.1530 |
| TOL_AvgTrlTime | 0.3801$^{\circ}$ | 0.3399$^{\circ}$ | -0.2016 | 0.1843 | 0.2374 | 0.1530 |
| VST_TotCor_Fw | **-0.4275\*** | -0.3956$^{\circ}$ | 0.1653 | -0.0183 | -0.0254 | -0.0191 |
| VST_Span_Fw | **-0.4147\*** | -0.3906$^{\circ}$ | 0.1290 | -0.0339 | -0.0265 | 0.0085 |
| VST_TotAvgTime_Fw | 0.0037 | -0.0366 | -0.1846 | 0.1010 | 0.1133 | 0.0479 |
| VST_TotCorAvgTime_Fw | -0.0101 | -0.0514 | -0.1887 | 0.1400 | 0.1322 | 0.0135 |
| VST_TotCor_Bw | -0.2895 | -0.2549 | 0.1689 | -0.1274 | -0.0815 | 0.0707 |
| VST_Span_Bw | -0.2859 | -0.2628 | 0.1160 | -0.0717 | -0.0245 | 0.0853 |
| VST_TotAvgTime_Bw | -0.0793 | -0.1233 | -0.1994 | -0.0228 | -0.0206 | -0.0001 |
| VST_TotCorAvgTime_Bw | -0.0548 | -0.1297 | -0.3422$^{\circ}$ | -0.0993 | -0.0226 | 0.1423 |

<sup>†</sup> Refers to the corrected scores.
<sup>ψ</sup> Refer to *Section 2.4* for the details of the neuropsychological variables listed in this table.
\* $p < 0.05$ (statistically significant at 5% level)

#### 3.3.3. Correlations of Age Measures with Impulsivity Variables

Correlations of chronological and brain age variables with BIS impulsivity measures for the AUD and CTL groups are shown in **Table 4**. A significant negative correlation between the non-planning score and brain age gap in the AUD group, suggesting that a higher brain age gap (i.e., higher acceleration of brain age) was associated with lower non-planning impulsivity. However, neither chronological age nor brain age showed any significant correlations with impulsivity variables. On the other hand, a negative correlation between the brain age gap and total impulsivity (r =-0.3805) was in the range of moderate effect size and can be considered for interpretation.

**Table 4.**
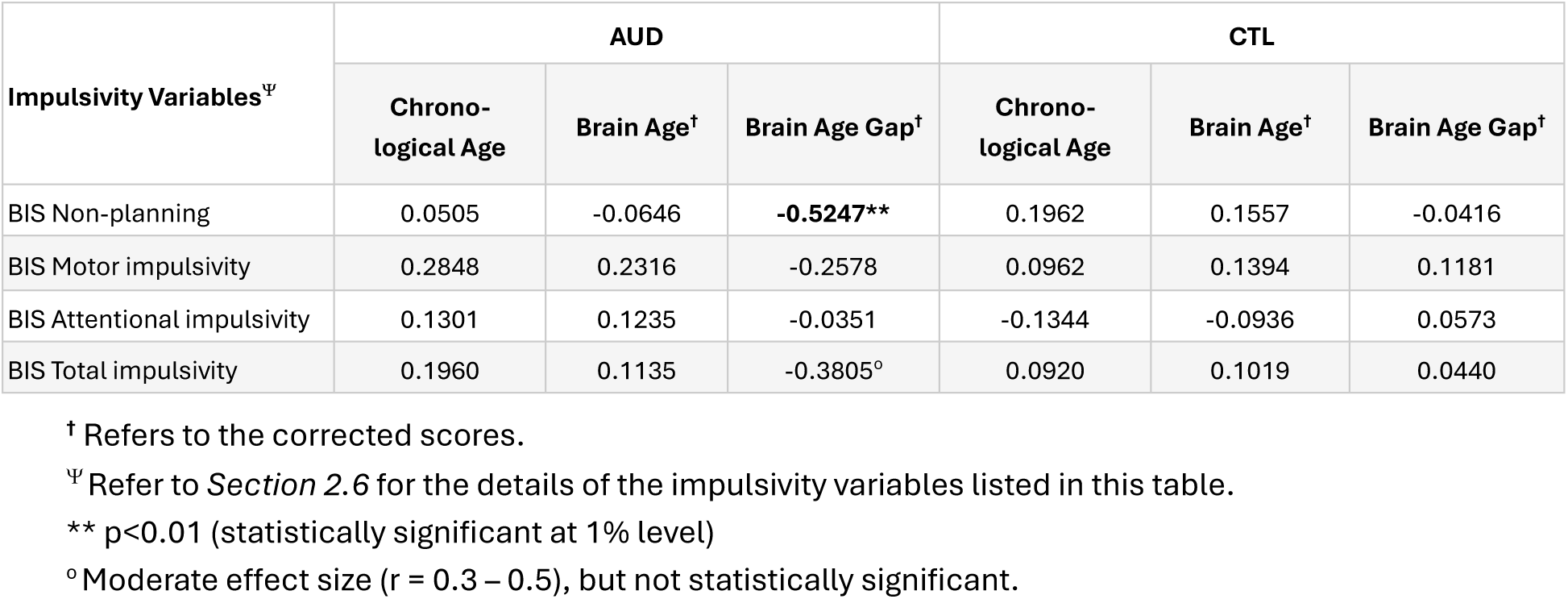
Correlations of specific age measures with impulsivity variables. Significant correlation is highlighted in bold font. Correlations with a moderate effect size (r = 0.3 – 0.5), but not statistically significant, have been marked with an omicron superscript (^ο^).

#### 3.3.4. Correlations of Age Measures with Alcohol Consumption Variables

The correlations of chronological and brain age variables with alcohol consumption measures for the AUD and CTL groups are shown in **Table 5**. A significant positive correlation was observed between the age of onset of regular drinking and the brain age gap in the AUD group. However, none of the other drinking measures showed any significant correlations with the age measures. On the other hand, some of the correlation coefficients, although not statistically significant, were in the range of moderate effect sizes and can be considered for interpretation: (a) age of onset of regular drinking was positively correlated with both chronological age (r = 0.3451) and brain age (r = 0.3160) in the CTL group; (b) number of drinking days per month during the regular drinking period was positively correlated with brain age in the AUD group (r = 0.3030) and negatively correlated with both chronological age (r =-0.3965) and brain age (r =-0.3228) in the CTL group.

**Table 5.**
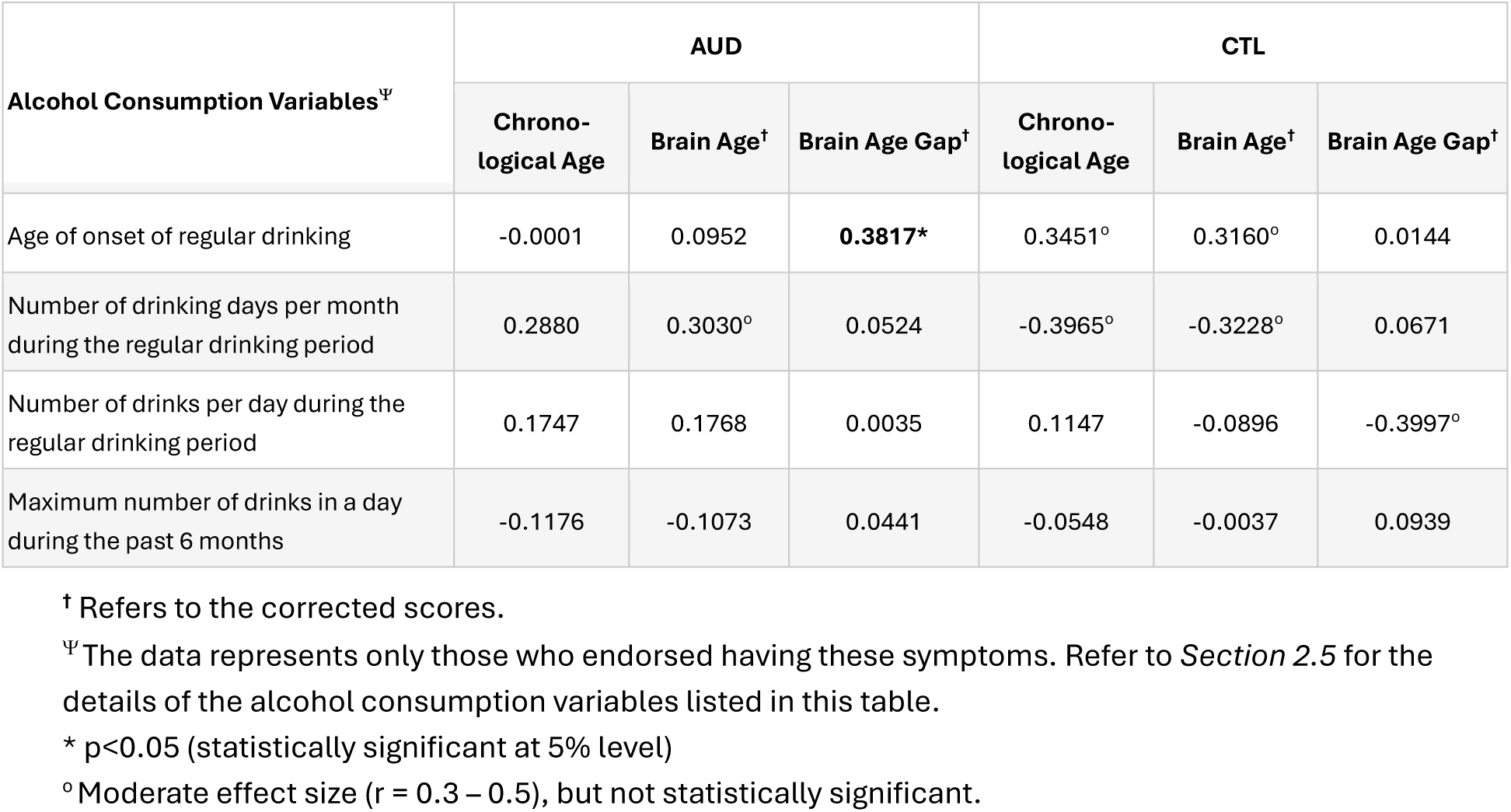
Correlations of specific age measures with alcohol consumption variables. Significant correlation is highlighted in bold font. Correlations with a moderate effect size (r = 0.3 – 0.5), but not statistically significant, have been marked with an omicron superscript (^ο^).

| Alcohol Consumption Variables<sup>Ψ</sup> | AUD |  |  | CTL |  |  |
| --- | --- | --- | --- | --- | --- | --- |
|  | Chrono-logical Age | Brain Age<sup>†</sup> | Brain Age Gap<sup>†</sup> | Chrono-logical Age | Brain Age<sup>†</sup> | Brain Age Gap<sup>†</sup> |
| Age of onset of regular drinking | -0.0001 | 0.0952 | **0.3817\*** | 0.3451$^{\circ}$ | 0.3160$^{\circ}$ | 0.0144 |
| Number of drinking days per month during the regular drinking period | 0.2880 | 0.3030$^{\circ}$ | 0.0524 | -0.3965$^{\circ}$ | -0.3228$^{\circ}$ | 0.0671 |
| Number of drinks per day during the regular drinking period | 0.1747 | 0.1768 | 0.0035 | 0.1147 | -0.0896 | -0.3997$^{\circ}$ |
| Maximum number of drinks in a day during the past 6 months | -0.1176 | -0.1073 | 0.0441 | -0.0548 | -0.0037 | 0.0939 |

<sup>†</sup> Refers to the corrected scores.
<sup>Ψ</sup> The data represents only those who endorsed having these symptoms. Refer to *Section 2.5* for the details of the alcohol consumption variables listed in this table.
\* $p < 0.05$ (statistically significant at 5% level)

## 4. Discussion

The objective of the present study was to predict brain age using neuroanatomical features such as cortical thickness and brain volume in individuals with a lifetime diagnosis of AUD. Results showed that AUD individuals showed an increase of 1.70 years in their brain age relative to their chronological age (i.e., brain age gap in the AUD group = 1.70 years), suggesting accelerated brain aging in these individuals. Brain age and brain age gap were found to be associated with some of the measures of neuropsychological performance, impulsivity, and alcohol consumption. The significance and implications of the major findings of the current study are detailed below.

### 4.1. Brain Age Measures between the AUD and CTL Groups

A vast majority of the AUD individuals, as illustrated in the distribution plot in **Figure 1**, consistently showed increased brain age. As shown in **Figure 2**, the corrected brain age for the AUD individuals was 1.70 years higher than their chronological age. Further, there was a significant correlation between chronological age and corrected brain age in AUD individuals (r = 0.9691; **Table 2**). In other words, the predicted brain age of AUD individuals was 1.70 years higher than their actual age. This finding indicates that AUD individuals, despite their current abstinence status, have accelerated brain aging, perhaps due to their history of AUD and chronic and/or heavy drinking. Previous neuroimaging studies have suggested that chronic alcohol use can lead to accelerated aging [19,90,91], as evidenced by anomalies in brain morphology as well as neurocognitive functions [19,92,93], which might further contribute to age-related dementia as an end outcome [19]. As the current study has used brain morphological features to estimate brain age, it is also worth mentioning that the same set of AUD participants had shown significantly smaller volumes in frontal lobe regions, such as left pars orbitalis, right medial orbitofrontal, right caudal middle frontal cortices, and in bilateral hippocampal regions in our previous study [32]. A previous study on brain age and AUD reported that both gray and white matter volume loss in the brain contribute to accelerated aging in individuals with chronic alcohol use [94].

While the decreased cortical thickness seen in abstinent AUD individuals could be associated with the severity of past alcohol misuse [95], there is evidence showing that age-related cognitive decline is further accelerated by chronic alcohol misuse [41,91,96].

Daily consumption of alcohol [97], including low-level alcohol consumption [98], was found to be associated with accelerated brain aging.

It may also be worthwhile to interpret the current findings in terms of other relevant domains of the brain and cognition. Studies have reported that brain structural abnormalities seen in AUD individuals are often associated with accompanying cognitive impairments [99–101], which can lead to accelerated cognitive aging and even dementia in chronic alcohol users [102–105]. A comparison of neuropsychological scores between the AUD and CTL groups has shown significantly poorer neurocognitive performance by the AUD individuals relative to the control group in both TOL and VST tests. This finding is further confirmed by our previous findings that the AUD participants showed relatively poorer performance in neuropsychological tests of problem-solving ability, visuospatial memory span, and working memory, compared to the healthy controls, in addition to changes in brain volume and white matter integrity [32]. Additionally, our past study also found that neuropsychological performance was correlated with several features of brain volume and white matter integrity [32], further validating the intricate relationship among brain structure, cognitive functioning, and brain aging.

### 4.2. Associations between Brain Age and Neuropsychological Performance

Correlational analysis (**Table 3**) revealed that increased brain age was significantly associated with poor neuropsychological performance related to executive functions such as planning and problem-solving in AUD individuals. In other words, AUD individuals with increased brain age showed poor performance on planning and problem-solving (as reflected by the number of trials taken by the individuals to solve the puzzles). This finding supports the view that brain age could be a reliable measure to estimate cognitive decline in various neurocognitive disorders, including in individuals diagnosed with AUD. On the other hand, in the visual memory task, only the chronological age showed a negative association with short-term memory performance (forward recall) in the AUD group, suggesting poor memory performance with advancing chronological age in individuals with AUD. It should also be noted that the correlation coefficients for the negative association between the short-term memory variables (total correct and span scores during the forward trials) and brain age were-0.3956 and-0.3906, respectively, in the AUD group, suggesting a moderate effect size [106] despite not reaching statistical significance, possibly suggesting poor memory performance associated with increased brain age. This could be due to lingering brain damage in individuals with a history of chronic AUD [28,107–109] and associated neurocognitive deficits [53], including memory problems [110,111].

However, these preliminary findings need confirmation from future studies with larger sample sizes.

Comparison of performance scores between the groups indicated that the AUD group showed significantly poorer neurocognitive performance in both the TOL and VST tests, relative to the CTL group (**Table 1**). These results support the previously reported findings on alcohol-related neuropsychological impairments in multiple domains, such as deficits in executive functioning, memory, and visuospatial processing [50–55,112]. A recent meta-analysis further confirmed that planning, problem-solving, and inhibitory abilities are significantly affected by alcohol misuse [40]. While the recovery of some cognitive processes is known to occur due to abstinence in AUD individuals [113,114], certain deficits can persist even after a prolonged abstinence [112]. Taken together, our findings suggest that measures of brain age can be useful markers of cognitive performance and brain health in AUD individuals, although these preliminary findings need further confirmation by larger studies in the future.

### 4.3. Associations between Brain Age and Impulsivity

Our findings showed that there was a significant negative correlation between the non-planning score and brain age gap in the AUD group, suggesting that those with lower non-planning impulsivity had higher brain age gap (i.e., accelerated brain age) (**Table 4**). Further, a negative correlation between the brain age gap and total impulsivity with a moderate effect size (r =-0.3805) was also observed in the AUD group, although this correlation did not achieve statistical significance due to the small sample size (N = 30). Although there are no previous studies on AUD that explored the impulsivity-brain age gap relationship, a recent study on depression reported that accelerated brain aging was associated with greater impulsivity and depression severity [115]. However, it should be noted that the associations between impulsivity and brain age gap are in the opposite direction in the depression study compared to our findings. This discord could be due to heterogeneity of both AUD and depressive disorder in terms of internalizing and externalizing traits, types of impulsivity, etiological pathways, and gender/sex differences [116–118]. For example, Boschloo et al. [117] showed that AUD was more strongly associated with high disinhibition, while thrill/adventure seeking was more strongly related to depression [117].

In terms of the neural underpinnings of impulsivity, previous studies have elucidated specific brain structures and neural networks associated with trait impulsivity [119–121]. It was also reported that anterior insula thickness and non-planning impulsivity showed a negative association with age, while there was a positive association between anterior insula thickness and non-planning impulsivity [122].

Our findings on group differences in impulsivity showed that individuals with AUD exhibited significantly increased impulsivity in all three categories and the total score (**Table 1**). However, interestingly, none of the impulsivity factors showed a significant association with actual age or brain age in either group, although previous studies have reported a negative correlation between chronological age and impulsivity [123,124]. It is possible that our small sample of males did not have the required age variations, especially the abstinent AUD group, most of whom were in their 40s with a mean age of 41.25 years.

Further, the vast majority of the AUD individuals were in abstinence from drinking after they received addiction treatment for their AUD diagnosis, which may have stabilized the variations in impulsive behaviors. Furthermore, the sample size (N = 30 per group) is rather too small to establish a significant association between age and impulsive traits; therefore, future studies should explore these associations using larger samples. Although we expected significant correlations between impulsivity and brain age in each group, we did not find this pattern in our results, except for a significant negative correlation of brain age gap with non-planning impulsivity (r =-0.5247) in AUD individuals. On the other hand, there was also a negative correlation of brain age gap and total impulsivity with a moderate effect size (r =-0.3806, but not significant), further emphasizing the impulsivity-brain age association in AUD individuals, although these preliminary findings warrant confirmation from future studies with larger sample sizes.

### 4.4. Associations between Brain Age and Alcohol Consumption

Our findings revealed that there was a significant positive association between the age of onset for regular drinking and the brain age gap in the AUD group, suggesting that those who started drinking at their later age showed accelerated brain aging. Further, there were several correlations with moderate effect sizes (greater than 0.3): (a) number of drinking days per month during the regular drinking period was positively correlated with brain age in the AUD group (r = 0.3030); (b) age of onset of regular drinking was positively correlated with both chronological age (r = 0.3451) and brain age (r = 0.3160) in the CTL group; (b) number of drinking days per month during the regular drinking period was negatively correlated with both chronological age (r =-0.3965) and brain age (r =-0.3228) in the CTL group. Although a lack of previous findings on these associations limits the interpretation of our current findings of correlations with moderate effect sizes across the measures of alcohol consumption and brain age, the quantity and frequency of alcohol consumption can impact brain age and hence brain health. However, as it is well-established that alcohol intake affects both the structure and function of the brain [125,126], it is understandable that brain age, a measure that depends on the integrity of brain structure and function, can be impacted by the measures of alcohol consumption.

Comparison of alcohol consumption measures across the groups revealed that the AUD group endorsed significantly increased alcohol consumption relative to the control group in all four drinking measures (**Table 1**). Given these differences, we expected stronger and widespread associations among alcohol consumption and brain age measures, especially in the AUD group. However, our findings on alcohol consumption and brain age measures are not very strong except for a single significant correlation between the age of onset for regular drinking and the brain age gap in the AUD group. Possible reasons for the lack of significant associations between alcohol consumption and brain age measures could be the following: First, the AUD individuals were maintaining abstinence from drinking at the time of clinical and MRI assessments, and the alcohol consumption measures reflect the past drinking history. Second, the sample size is too small to establish stronger and significant associations between the drinking measures and brain age. Further studies with larger sample sizes using subgroups of AUD individuals at different stages or severity of past and current AUD diagnoses are needed to ascertain the relationship between brain age and drinking measures.

### 4.5. Limitations of the Current Study and Suggestions for Future Research

While the study has yielded some interesting findings, several limitations need to be acknowledged. First, the sample size was relatively small for the predictive models of machine learning; therefore, the validity of the prediction, as well as the generalizability of the findings, is rather limited in scope. Second, the study groups were not matched for age, as the control participants were significantly younger than their AUD counterparts. Finally, the sample comprised only male participants, and therefore, generalizability is limited.

Future studies should use large samples of AUD individuals, preferably matched for age, sex, and other important sample characteristics, to obtain realistic brain measures.

Further, the association among brain age, neural features, and behavioral/cognitive features may be examined by future studies using samples from different age cohorts (e.g., children, adolescents, young adults, etc.) to understand potential etiological mechanisms underlying alcohol use and related disorders across brain development. Future research on brain age in AUD may also consider using other important structural and functional brain features, including brain connectivity measures, as done by some of the recent brain age studies on other neuropsychiatric disorders. It may be interesting and important to compare various prediction algorithms of both machine learning and linear models to ascertain and validate the findings across multiple studies. Lastly, future studies may also compute brain age using measures from various domains such as neuroimaging (i.e., structural and functional MRI measures, resting and task-related EEG measures) and neurocognition (e.g., neuropsychological scores), and examine their associations with measures of personality traits (e.g., internalizing and externalizing characteristics), genomics (e.g., polygenic scores), and environment (e.g., SES, family, peers, etc.).

## 5. Conclusions

Brain age measures are assuming greater importance in predicting cognitive decline and accelerated biological aging in various disorders, including substance use disorders. The current study found that the AUD group showed an increase of 1.70 in their brain age relative to their chronological age, suggesting accelerated brain aging and cognitive decline/deficit in these individuals. In the AUD group, higher brain age was significantly associated with poor executive functioning, while a larger brain age gap was associated with lower non-planning impulsivity and later age of onset for regular drinking. Further, although statistically not significant, there were several other correlations with moderate effect sizes across brain age measures and the measures of neurocognition, impulsivity, and alcohol consumption, which should be confirmed by future studies with larger sample sizes. Our major finding that AUD individuals, despite maintaining abstinence, manifested accelerated brain aging can be explained by potential brain damage caused by the history of chronic alcohol use. It is suggested that future studies should use multimodal neural measures to compute brain age and also explore associations between brain age measures and other measures of personality, genomics, and environment.

## Author Contributions

Conceptualization: C.K., B.P., B.A.A., A.K.P.; Methodology: C.K., B.P., B.A.A., A.K.P., S.K.; Data Collection: B.A.A.; Data Curation: B.A.A., C.K.; Formal Analysis: C.K.; Manuscript Preparation: C.K., B.A.A., B.P.; Review and Editing: B.P., B.A.A., A.K.P., D.B.C., S.K., G.P., J.L.M., S.S.D., C.R., A.B., W.K.; Funding Acquisition: B.P., C.K., J.L.M., A.K.P. All authors have read and agreed to the published version of the manuscript.

## Institutional Review Board Statement

The study was approved by the Institutional Review Boards of all three research sites of the NYS state, i.e., SUNY Downstate Health Sciences University, Nathan Kline Institute for Psychiatric Research, and New York University [NYU Study # i16-02192 (Brain Dysfunction and Alcoholism), 10 March 2017, and NYU Study # i21-01219 (Brain Function and Neurogenomic Influences on AUD Risk and Resilience), 21 March 2022].

## Informed Consent Statement

Informed consent was obtained from all subjects involved in the study.

## Data Availability Statement

The data presented in this study will be made available to the researchers upon request to the corresponding author.

## Acknowledgments

In memory of Henri Begleiter, founder and longtime mentor of the Neurodynamics Laboratory, we acknowledge with great admiration his seminal scientific contributions to the field. We are sincerely indebted to his charismatic leadership and luminous guidance, truly inspired by his scientific mission and vision, and highly motivated to carry forward the work he fondly cherished. We are grateful for the valuable technical assistance of Carlene Haynes, Joyce Alonzia, Chamion Thomas, Alec Musial, Kristina Horne, Talia Stern, and Abigail Freeman.

## Abbreviations

Thefollowing abbreviations are used in this manuscript
AUD: Alcohol Use Disorder
BIS: Barratt Impulsiveness Scale
GM: Gray Matter
MAE: Mean Absolute Error
MMSE: Mini-Mental State Examination
MPRAGE: magnetization-prepared rapid gradient-echo
MRI: Magnetic Resonance Imaging
RMSE: Root Mean Squared Error
SSAGA: Semi-Structured Assessment for the Genetics of Alcoholism
TOL: Tower of London Test
VST: Visual Span Test
WM: White Matter

## Appendix

**Table A1.**
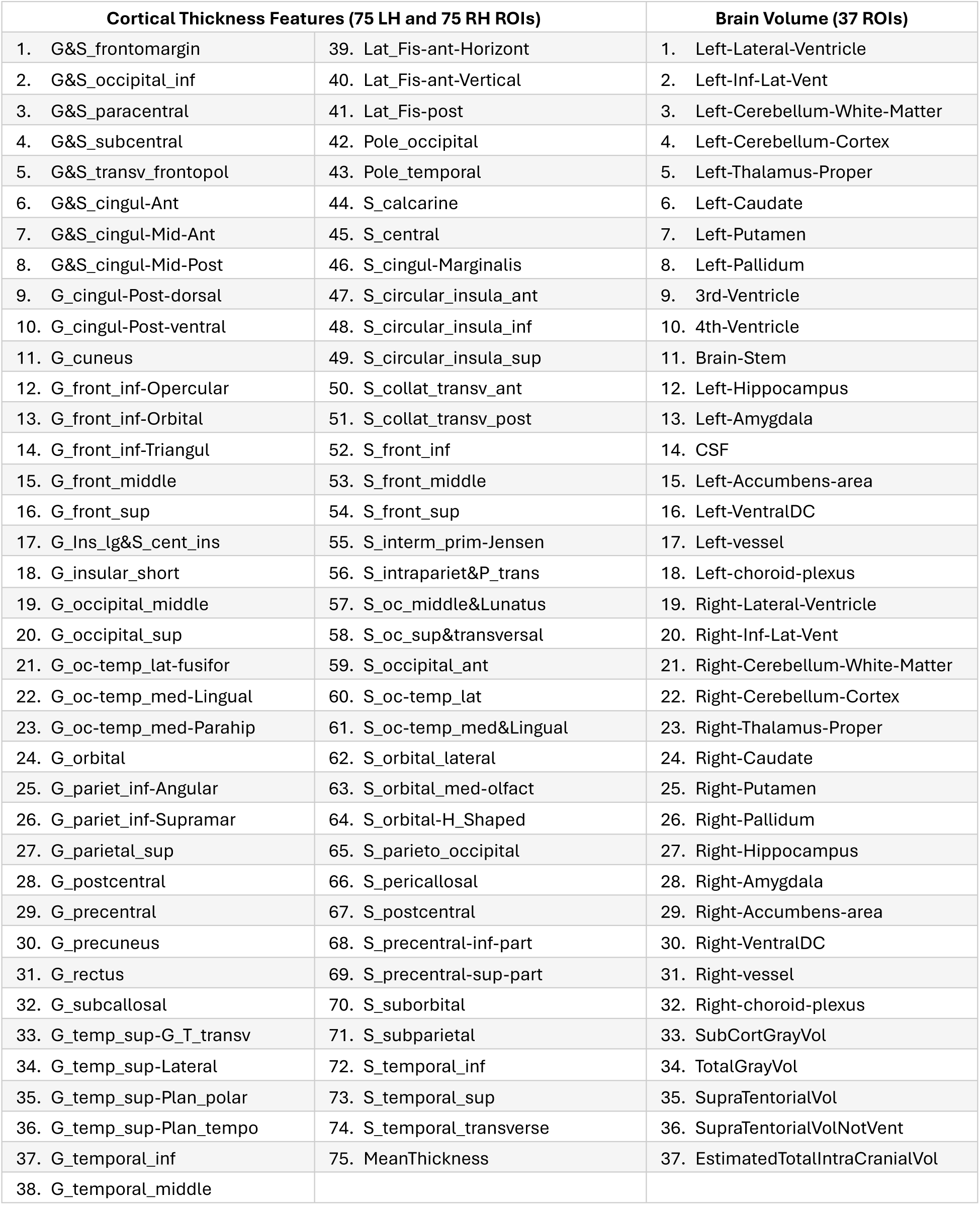
List of 187 anatomical features used in the prediction of brain age.

| Cortical Thickness Features (75 LH and 75 RH ROIs) |  | Brain Volume (37 ROIs) |
| --- | --- | --- |
| 1. G&S_frontomargin | 39. Lat_Fis-ant-Horizont | 1. Left-Lateral-Ventricle |
| 2. G&S_occipital_inf | 40. Lat_Fis-ant-Vertical | 2. Left-Inf-Lat-Vent |
| 3. G&S_paracentral | 41. Lat_Fis-post | 3. Left-Cerebellum-White-Matter |
| 4. G&S_subcentral | 42. Pole_occipital | 4. Left-Cerebellum-Cortex |
| 5. G&S_transv_frontopol | 43. Pole_temporal | 5. Left-Thalamus-Proper |
| 6. G&S_cingul-Ant | 44. S_calcarine | 6. Left-Caudate |
| 7. G&S_cingul-Mid-Ant | 45. S_central | 7. Left-Putamen |
| 8. G&S_cingul-Mid-Post | 46. S_cingul-Marginalis | 8. Left-Pallidum |
| 9. G_cingul-Post-dorsal | 47. S_circular_insula_ant | 9. 3rd-Ventricle |
| 10. G_cingul-Post-ventral | 48. S_circular_insula_inf | 10. 4th-Ventricle |
| 11. G_cuneus | 49. S_circular_insula_sup | 11. Brain-Stem |
| 12. G_front_inf-Opercular | 50. S_collat_transv_ant | 12. Left-Hippocampus |
| 13. G_front_inf-Orbital | 51. S_collat_transv_post | 13. Left-Amygdala |
| 14. G_front_inf-Triangul | 52. S_front_inf | 14. CSF |
| 15. G_front_middle | 53. S_front_middle | 15. Left-Accumbens-area |
| 16. G_front_sup | 54. S_front_sup | 16. Left-VentralDC |
| 17. G_Ins_lg&S_cent_ins | 55. S_interm_prim-Jensen | 17. Left-vessel |
| 18. G_insular_short | 56. S_intrapariet&P_trans | 18. Left-choroid-plexus |
| 19. G_occipital_middle | 57. S_oc_middle&Lunatus | 19. Right-Lateral-Ventricle |
| 20. G_occipital_sup | 58. S_oc_sup&transversal | 20. Right-Inf-Lat-Vent |
| 21. G_oc-temp_lat-fusifor | 59. S_occipital_ant | 21. Right-Cerebellum-White-Matter |
| 22. G_oc-temp_med-Lingual | 60. S_oc-temp_lat | 22. Right-Cerebellum-Cortex |
| 23. G_oc-temp_med-Parahip | 61. S_oc-temp_med&Lingual | 23. Right-Thalamus-Proper |
| 24. G_orbital | 62. S_orbital_lateral | 24. Right-Caudate |
| 25. G_pariet_inf-Angular | 63. S_orbital_med-olfact | 25. Right-Putamen |
| 26. G_pariet_inf-Supramar | 64. S_orbital-H_Shaped | 26. Right-Pallidum |
| 27. G_parietal_sup | 65. S_parieto_occipital | 27. Right-Hippocampus |
| 28. G_postcentral | 66. S_pericallosal | 28. Right-Amygdala |
| 29. G_precentral | 67. S_postcentral | 29. Right-Accumbens-area |
| 30. G_precuneus | 68. S_precentral-inf-part | 30. Right-VentralDC |
| 31. G_rectus | 69. S_precentral-sup-part | 31. Right-vessel |
| 32. G_subcallosal | 70. S_suborbital | 32. Right-choroid-plexus |
| 33. G_temp_sup-G_T_transv | 71. S_subparietal | 33. SubCortGrayVol |
| 34. G_temp_sup-Lateral | 72. S_temporal_inf | 34. TotalGrayVol |
| 35. G_temp_sup-Plan_polar | 73. S_temporal_sup | 35. SupraTentorialVol |
| 36. G_temp_sup-Plan_tempo | 74. S_temporal_transverse | 36. SupraTentorialVolNotVent |
| 37. G_temporal_inf | 75. MeanThickness | 37. EstimatedTotalIntraCranialVol |
| 38. G_temporal_middle |  |  |

